# Marine transmissible cancer navigates urbanised waters, threatening to spillover

**DOI:** 10.1101/2023.04.14.536605

**Authors:** M. Hammel, F. Touchard, E. A. V. Burioli, L. Paradis, F. Cerqueira, E. Chailler, I. Bernard, H. Cochet, A. Simon, F. Thomas, D. Destoumieux-Garzón, G. M. Charrière, N. Bierne

## Abstract

Inter-individual transmission of cancer cells represents a unique form of microparasites increasingly reported in marine bivalves. In this study, we sought to understand the ecology of the propagation of *Mytilus trossulus* Bivalve Transmissible Neoplasia 2 (MtrBTN2), a transmissible cancer affecting four *Mytilus* mussel species worldwide. We investigated the prevalence of MtrBTN2 in the mosaic hybrid zone of *M. edulis* and *M. galloprovincialis* along the French Atlantic coast, sampling contrasting natural and anthropogenic habitats. We observed a similar prevalence in both species, likely due to the spatial proximity of the two species in this region. Our results showed that ports had higher prevalence of MtrBTN2, with a possible hotspot observed at a shuttle landing dock. No cancer was found in natural beds except for two sites close to the hotspot, suggesting spillover. Ports may provide favourable conditions for the transmission of MtrBTN2, such as high mussel density, stressful conditions, sheltered and confined shores, or buffered temperatures. Ships may also spread the disease through biofouling. Our results suggest ports may serve as epidemiological hubs, with maritime routes providing artificial gateways for MtrBTN2 propagation. This highlights the importance of preventing biofouling on docks and ship hulls to limit the spread of marine pathogens hosted by fouling species.

## 1. Introduction

Transmissible cancers are caused by malignant cell lineages that have acquired the ability to infect new hosts through the transmission of living cancer cells. Fourteen transmissible cancer lineages have been described to date: one in dogs (canine transmissible venereal tumour, CTVT [1, 2]), two in Tasmanian devils (devil facial tumour, DFT1 and DFT2 [3, 4]), and eleven in different marine bivalve species (bivalve transmissible neoplasia, BTNs [5–10]). While direct contact is necessary for CTVT and DFTs transmission, via coitus for the former and biting for the latter, the transmission of BTNs between shellfish is assumed to occur through the seawater column [11, 12]. *Mytilus* spp. mussels are affected by two BTN lineages known as MtrBTN1 and MtrBTN2 that originate from two different *M. trossulus* founder hosts. MtrBTN2 is distributed worldwide and has crossed the species barrier several times to infect four *Mytilus* species: *M. trossulus* in East Asia and Northern Europe [13–15], *M. chilensis* in South America [9], and *M. edulis* and *M. galloprovincialis* in Europe [9, 16]. Cancerous process in mussels was firstly described fifty years ago as disseminated neoplasia (DN, [17]) but the demonstration that some DNs are transmissible is recent [6, 7]. As non-transmissible neoplasia also occurs in mussels [9, 16], a genetic diagnosis is required to ensure the identification of MtrBTNs. The oldest known genetically validated MtrBTN2 was sampled in 2009 [9]. Genetic data suggest that it could be at least one or two orders of magnitude more ancient, although dating is difficult due to an apparent acceleration of the mitochondrial clock [15, 16].

The global distribution of the MtrBTN2 lineage remains enigmatic [9, 14]. The other known transmissible cancer with such a worldwide distribution is CTVT in dogs. CTVT emerged 4,000 to 8,000 years ago, probably in Eastern Asia, and has expanded rapidly worldwide over the past 500 years, probably facilitated by the development of maritime transportation [18]. Transmission of MtrBTN2 within mussel populations occurs presumably through filter-feeding (discussed in [11, 19]). MtrBTN2 cells can survive for several days in seawater [11], which greatly increases their chances of infecting new hosts. However, it is unlikely that transport of cancer cells by marine currents alone explains such a global distribution. Indeed, the dilution of cancer cells in the marine environment may strongly limit the success of transmission over long distances. Shipping traffic has been proposed as the most likely explanation for the global distribution of MtrBTN2 -i.e., the transport of disease-carrying mussels on ship hulls [9, 14, 15]. To our knowledge, these claims have remained speculative and no formal evidence of the effect of shipping traffic and port habitat on the epidemiology of BTN has been provided to date.

European mussels are affected by MtrBTN2, with a much higher prevalence in *M. edulis* (of the order of 1/100) than in *M. galloprovincialis* (of the order of 1/1000) [16]. These two host species are parapatric in Europe and can coexist in contact zones where hybridisation is taking place [20–22]. Associations between genetic backgrounds and environmental variables indicate that the environment partly influences the structure and maintenance of the hybrid zones: in the study area, *M. edulis* genotypes are more frequent in sheltered habitats under freshwater influence, while *M. galloprovincialis* genotypes are more frequent in habitats exposed to wave action and ocean salinities [23, 24]. In this context, the distribution and propagation of MtrBTN2 is likely to be influenced by host genetic background and population composition [16], as well as by other environmental factors such as population density, reduced hydrodynamics, and stressful environmental factors (e.g., heat, salinity or pollution stresses). MtrBTN2 global distribution indicates its resilience across a broad range of temperature with varying seasonal patterns [14], reflecting host species distribution range. Prior to evidence of horizontal transmission, pollutants have been suggested as a potential cause of DN; however, the association with disease prevalence remained inconclusive [25–27], and as we now know that some neoplasias are transmissible, it is no longer the initial carcinogenesis but the transmission that needs to be explained. Seasonal influence on DN prevalence was observed at a local scale (higher during winter in *M. trossulus*, [28, 29], and *M. edulis*, [30]), whereas no significant influence of seasonality was noted at the European scale [31]. Unfortunately, in the absence of genetic diagnostics, we cannot know how many of these DNs were indeed BTNs (i.e. transmissible) or not, and rigorous evaluation of MtrBTN2 prevalence seasonality is still lacking. In brief, little is known about the ecology of MtrBTN2 in relation to its host and the possible environmental factors influencing the epidemiology of this transmissible cancer.

In the present study, we investigated the occurrence of MtrBTN2 in a survey area in Southern Brittany (France) which has contrasting natural and anthropogenic habitats and a mosaic distribution of *M. edulis* and *M. galloprovincialis*. We sampled 40 natural populations as well as 9 mussel farms, 7 floating buoys, and 20 ports. To detect MtrBTN2-infected mussels, we used digital PCR and real-time PCR, amplifying one nuclear and two mitochondrial genes with published and newly designed primers targeting specific variants. We found a low prevalence of MtrBTN2 (23/1516), equally affecting the two coexisting *Mytilus* species. Interestingly, most affected sites (7/9) were ports, suggesting that ports could be favourable habitats for MtrBTN2 transmission and/or that maritime transport could play a role as a vector for the spread of the disease. This highlights the possible role of ports as epidemiological hubs for MtrBTN2 propagation.

## 2. Material and methods

### Sampling

We collected 1516 *Mytilus* spp. mussels from 76 sites of the French Atlantic coast, from the Bay of Quiberon to Pornic, between January and March 2020. Approximately 20 mussels were sampled at each site (Table S1), allowing us to have a large number of closely located sites. We collected hemolymph from adductor muscle (with a 1 ml syringe, 26G needle) and a piece of mantle; both samples were fixed with 96% ethanol as described in [16]. DNA extraction was performed with the NucleoMag 96 Tissue kit (Macherey-Nagel) using a Kingfisher Flex extraction robot (serial number 711-920, ThermoFisher Scientific). We followed the kit protocol with a modified lysis time of 1 hour at 56°C and modified the volumes of some reagents: beads were diluted twice in water, 200 µl of MB3 and MB4, 300 µl of MB5 and 100 µl of MB6. DNA concentration (ng/μl) was quantified using the Nanodrop 8000 spectrophotometer (ThermoFisher Scientific). DNA from hemolymph was used for MtrBTN2 detection and DNA from mantle was used for mussel genotyping (Figure S1).

### MtrBTN2 detection

As the prevalence of MtrBTN2 is expected to be low [16, 32] and the number of samples is high (n = 1516), we chose to pre-screen by pooling 12 samples (pooled screening step; Figure S1) and then demultiplexing positive pools to specifically target cancerous samples (simplex screening step; Figure S1). Prior to pooling, DNA concentrations were adjusted to 10 ng/µl to obtain an unbiased representation of each sample in the pool. This allowed us to considerably reduce the cost and time of detection, but we acknowledge that this protocol could result in some very early stages of the disease being missed. To partly circumvent this problem, MtrBTN2 detection in pools was performed by digital PCR (ddPCR) targeting one nuclear (Elongation Factor, EF) and one mitochondrial (Control region, described in [9]) marker (Table S2). ddPCR is a sensitive method, based on the Taqman method which requires two primers and one probe, and which directly estimates the copy number of the targeted sequence. In addition, the use of a mitochondrial marker should provide a more sensitive detection as the mitochondrial genome has more copies than the nuclear genome. ddPCR primers design and analysis was subcontracted to the company IAGE using the QIAcuity™ system (QIAGEN). Positive pools were then demultiplexed (simplex screening step; Figure S1) and MtrBTN2 detection was performed by real-time PCR targeting one nuclear (Elongation Factor, EF1α-i3, described in [11]) and one mitochondrial (cytochrome c oxidase I, mtCOI-sub) marker (Table S2). To design the mtCOI- sub primers, we used sequences of *M. edulis*, *M. galloprovincialis*, *M. trossulus*, and MtrBTN2 available from the National Center for Biotechnology Information (NCBI). Please note that mtCOI-sub primers were initially designed to amplify a polymorphism that discriminates between two MtrBTN2 sub-lineages “a” and “b” [16]. However, attempts to identify sub- lineages based on melting temperature (Tm) did not yield conclusive results, most likely due to additional polymorphism. Amplification of EF1α-i3 and mtCOI-sub markers was performed using a three-step cycling protocol (95°C for 10 s, 58°C for 20 s, 72°C for 25 s) for 40 cycles. We carried out real-time PCRs using the SensiFAST^TM^ SYBR No-ROX Kit (Bioline) and the LightCycler 480 Real-Time PCR System (Roche Diagnostics). We also confirmed the positive samples diagnosed by real-time PCR using ddPCR (simplex screening step; Figure S1). The specificity of both methods and markers was tested using various control samples of *M. edulis*, *M. galloprovincialis*, *M. trossulus*, and MtrBTN2 samples of early, moderate, or late stages (based on cytological observations, following [32], Tableau S3). To test the sensitivity, we used positive controls with MtrBTN2 DNA diluted 10-fold (x10) and 100-fold (x100) in *M. edulis* DNA.

Real-time PCR results were analysed using the LightCycler480 software. Two parameters were extracted: threshold cycle (CT), obtained by Absolute Quantification analysis using the Second Derivative Maximum method, and Tm, obtained by Tm calling analysis. Positive samples were defined as those with a CT of less than 35 cycles and a Tm between 77.46 to 78.71 (based on Tm values of the positive control samples; see result, Figure S2). ddPCR results were analysed with the QuantaSoft™ Analysis Pro software (Bio-Rad), which provides the copy number for each sample. Samples were considered positive when there were more than two positive droplets.

### Mussel genotyping

We used 10 biallelic SNPs known to discriminate between *M. edulis* and *M. galloprovincialis* mussel species (Table S4). These markers were identified as being ancestry-informative (fixed for alternative alleles in *M. edulis* and *M. galloprovincialis*) in the [33] dataset and were subsequently confirmed as near diagnostic by analysis of larger datasets [22, 34]. Genotyping was performed using the Kompetitive Allele Specific PCR (KASP) method [35]. As we wanted to know the mean ancestry of each sample (G ancestry) and reduce the cost of genotyping, we developed a multiSNPs marker by multiplexing the 10 SNPs. Rapidly, 1µL of assay mix (KASP-TF V4.0 2X Master Mix, 1X target concentration; primers, 1µM target concentration; HyClone™ HyPure water) and 0.5 µL of DNA at 10 ng/µL were mixed in qPCR 384-well plates using Labcyte Echo525 (BECKMAN COULTER). KASP analysis was performed on the LightCycler 480 Instrument (Roche Diagnostics) with the following thermal cycling conditions: initialisation 15 min at 95°C, first amplification 20 s at 94°C and 1 min at 61°C to 55°C with steps of 0.6°C for 10 cycles, second amplification 20 s 94°C and 1 min at 55°C for 29 cycles, read 1 min at 37°C and 1 s at 37°C.

KASP results are a combination of two fluorescence values, one for allele X, and another for allele Y. Here, we oriented all SNPs so that allele X corresponds to *M. edulis* and allele Y to *M. galloprovincialis.* Following [36], we transformed the data to obtain a single measure of the relative fluorescence of the two alleles, using the following formula: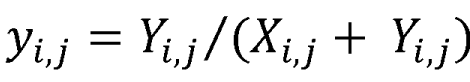; where X_i,j_ and Y_i,j_ are the fluorescence values of allele X and allele Y, respectively for, i^th^ SNP of the j^th^ sample. To scale y_i,j_ value from 0 when *M. edulis* allele fluorescence dominates at the 10 SNPs to 1 when *M. galloprovincialis* allele fluorescence dominates, we used the following formula: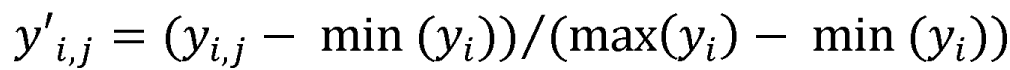; where y_i,j_ is the relative fluorescence value for the i^th^ SNP of the j^th^ sample. In the results section and figures, 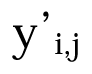corresponds to “G ancestry”.

### Environmental data

#### Source of environmental data

The 76 sampling sites were selected to cover four different habitats (ports, natural beds, farms and floating buoys) presenting contrasted environments and genetic backgrounds (*M. edulis*, *M. galloprovincialis* and hybrids). The description of environmental data considered in this study is reported in Table S5 and all site information is available in Table S1. Note that natural beds are intertidal sites, while the other habitats are subtidal. During sampling, we collected two habitat discrete descriptive variables: population density (1: isolated, 2: patchy beds, 3: continuous beds) and wave action (1: exposed, 2: sheltered). We used the E.U. Copernicus Marine Service Information (doi: 10.48670/moi-00027) to access temperature (°C), salinity (PSU 1e-3) and surface current velocity (m.s-1) data for each site (monthly recorded from 2021, resolution of 0.028°x0.028°). The velocity variable was formed by combining its Northward (N) and Eastward components, as follow: 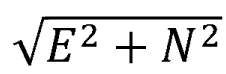We also used EMODnet Human Activities (source: EMSA Route Density Map) to retrieve the maritime traffic density of fishing, passenger, and recreational boats in the study area (seasonal record from 2018, 2019 and 2020, resolution 1x1 km).

#### Preparation of environmental data

Among the 76 sampled sites, 13 fell outside the coverage scope of the Copernicus model (i.e situated beyond the model’s pixel boundaries). Consequently, for these sites (see Table S1), we were unable to extract temperature, salinity, or velocity data. Certain sites recorded a velocity of 0 for all month (Table S1), raising concerns about the model resolution for this variable. To validate the accuracy of Copernicus data, we computed the mean values of neighbouring pixels (MNP, within a 0.05° buffer zone) and compared them to the pixel values at the respective sites of interest (SP, referred to as the ’central’ pixel, Table S6). We note that, due to the proximity of our sampled sites to the coastline, the number of neighbouring pixels varied within a certain range (Table S6). Notably, there was no correlation between these variations and the absolute difference between SP and MNP values (Figure S3C). The comparison between SP and MNP values was carried out for all sites, encompassing temperature, salinity, and surface current velocity, and the results are presented in Figure S3. We observed a robust positive correlation for temperature and salinity variables, providing confidence in the reliability of the extracted data for our small-scale analysis. However, as expected, the inconclusive correlation for surface current velocity led us to exclude this environmental variable from the analyses.

In the statistical analysis (see section 2.5) we used the annual average of salinity and boat density (maritime traffic). For temperature, we used the annual variance as we identified strong seasonal variations in temperature that would be misinterpreted if only the annual average temperature were used (Figure S4).

### Statistical analysis: relationship between MtrBTN2 prevalence, host genotype and environmental variables

To ascertain whether cancer prevalence varied among the four habitats sampled (ports, natural beds, buoys and farms), we performed Fisher’s exact test on the contingency table (*fisher.test()* function of the *stats* package *v4.1.3* [37]). Subsequently, we aimed to determine whether specific environmental variables could provide a more comprehensive explanation of MtrBTN2 prevalence across these habitats. To do so, we pursued the following steps: an exploratory analysis with a Principal Component Analysis (PCA), an examination of data dependencies (Spearman correlation test and Moran’s I test), and lastly, a regression model adapted to our dataset (spatial logistic regression model). All statistical analyses were performed in R *v4.1.3* [37].

#### Principal component analysis

As an exploratory analysis, we firstly performed a PCA to investigate the relationship between the quantitative environmental variables. Missing values were estimated using *imputePCA()* (*missMDA* package *v1.18* [38]), prior to performing the PCA using *PCA()* (*FactoMineR* package *v2.6* [39]).

#### Spearman correlation test

To test for correlation between environmental variables, we performed a non-parametric test of Spearman using *rcorr()* function from *Hmisc* package [40], that automatically excludes missing data.

#### Moran I test for spatial autocorrelation

Given the proximity of the sampled sites, which constitutes a fine spatial matrix, we used the Moran’s I statistic to test for spatial autocorrelation among the environmental variables. After converting longitude and latitude data into UTM coordinates (*spTransform()* function of *rgdal* package v1.6-7 [41]), this analysis was performed using the moran.test() function (*spdep* package version 1.2-8 [42]) with the k-nearest neighbour classification.

#### Spatial logistic regression model

The association of MtrBTN2 prevalence and environmental variables was tested at the site level using a multivariate spatial logistic regression (*fitme()* function, *spaMM* package *v4.3.0* [43]). Spatial autocorrelation was accounted for by including a Matern covariance function as a random effect (i.e. *Matern(1 | Longitude +Latitude)*). MtrBTN2 prevalence was represented as a binary matrix (*N*, *N_site_*-*N*) with *N* and *N_site_* indicating site-specific MtrBTN2 case counts and total number of sampled mussels, respectively. The following explanatory variables were included into the full model: temperature variance, salinity mean, population abundance, wave action, average *M. galloprovincialis* ancestry of the mussel host population (G ancestry), maritime traffic (as described above, section 2.4). We also included a binary variable "port" into the model, where 1 denotes port habitat and 0 represents other habitats. Model suitability was evaluated using the residual diagnostics tool implemented in the “DHARMa” package [44]. To assess the significance of fixed effects (environmental variables), we performed Bartlett- corrected likelihood ratio tests (LR tests) by comparing the full model with a model that excluded the variable of interest (*fixedLRT()* function of *spaMM* package *v4.3.0* [45]). To account for bias associated with rare events and collinearity of environmental variables (see results), LR tests were performed using 999 bootstraps.

## 3. Results

### Validation of the screening method

In order to detect MtrBTN2 in thousands of individuals, we developed dedicated ddPCR and qPCR screening tools. The design of real-time PCR and ddPCR primers and probes allowed us to amplify nuclear and mitochondrial markers specific to *M. trossulus* mussels and MtrBTN2 (Figure S5, TableS7). However, for the mtCOI-sub locus, the melting temperature (Tm) range of MtrBTN2 was different from *M. trossulus* controls and allowed us to specifically detect MtrBTN2 based on Tm values (Figure S2). With ddPCR, all MtrBTN2 control samples were detected, regardless of marker or dilution, except for one MtrBTN2 sample diluted at x100 which was negative for the EF locus (Figure S5). Copy numbers appeared higher for the mtCR marker, showing a higher sensitivity for this mitochondrial marker. The same pattern was observed for EF1α-i3 and mtCOI-sub in real-time PCR, with a higher sensitivity of the mitochondrial marker based on the lower CT values observed (Figure S5). However, two MtrBTN2 samples diluted at x100 were negative for mtCOI-sub and positive for EF1α-i3. Even though the mitochondrial marker appears to be more sensitive in both methods, we chose to screen for MtrBTN2 using both markers in order to have comparisons and additional validation.

### MtrBTN2 screening reveals a low prevalence

We amplified nuclear and mitochondrial markers using ddPCR (EF, mtCR) and real-time PCR (EF1α-i3, mtCOI-sub) to detect the presence of MtrBTN2 in the 1516 mussels sampled from the 76 sites of the study area (Figure S1). All results of the pooled and simplex screening are presented in TableS8.

The pooled screening step using ddPCR revealed 25/127 positive pools: 9/127 with EF, 14/127 with mtCR, and 2/127 were positive for both markers (Figure S5). As the 14 mtCR-positive pools showed low copy numbers (median = 7.59), we assume that EF did not amplify because of a detection threshold. Although the 9 EF-positive pools were not expected, they were included for further investigation by simplex screening (see below). For pools positive for both markers, the EF copy numbers appear relatively low compared to mtCR copy numbers. This is due to the high mitochondrial fluorescence masking the nuclear fluorescence and underestimating EF copy numbers. To eliminate this bias in the simplex ddPCR step (Figure S1), we amplified the two markers separately. In order to maximise cancer detection in the simplex screening step, we chose to keep the pools positive for at least one gene, corresponding to 25 positive pools of 12 samples (n = 300).

We performed the simplex screening step using real-time PCR and then confirmed positive samples with ddPCR (Figure S1). In total, 44/300 samples were positive for at least one real-time PCR marker: 16 with EF1α-i3 only, 23 with mtCOI-sub only, and 5 with both markers (Figure S5, Table S9). Pooled and simplex screening results are mostly consistent, as 20 of the 25 positive pools had at least one individual detected by real-time PCR amplification (Table S10). The 5 pools without positive samples had low ddPCR copy numbers (less than 3.37 for EF and less than 6.65 for mtCR). These false positives were to be expected as we intentionally applied low threshold values in the pooled screening step. Two samples from mtCR-positive pools were found to be positive for both markers at the simplex level, which was expected as the mitochondrial marker is likely to be more sensitive than the nuclear marker. On the contrary, we did not expect to observe samples positive for EF1α-i3/EF and negative for mtCOI-sub/mtCR. However, all but one of the individuals from the EF-positive pools were positive only for the EF1α-i3 marker (Table S10). The absence of mitochondrial genes in these samples does not seem to be related to a detection threshold issue, especially as ddPCR and real-time PCR target two different genes (mtCR and mtCOI-sub). We suspect that some host alleles that were present at low frequencies in the ‘dock mussels’ populations (the hybrid lineage found in the port of Saint-Nazaire [34]) may sometimes be amplified with EF1α-i3. This could be explained by shared polymorphism between host species rather than the presence of MtrBTN2. Therefore, the 16 samples positive only for EF1α-i3 were conservatively considered as false positives (note they were mostly sampled in the port of Saint-Nazaire, which does not affect our main result). For the remaining 28 samples, we performed ddPCR with both markers to confirm the real-time mitochondrial PCR diagnosis (Figure S1, TableS9). For 23/28 samples, mtCR ddPCR confirmed the mtCOI-sub real-time PCR diagnosis. For 10/23 samples positive only for mtCOI-sub, ddPCR revealed few EF copies. Finally, we considered all samples positive for mitochondrial markers with both methods to be cancerous samples. We found 23/1516 positive samples from 9/76 sites in which the number of cases varied from 1 to 8 (Figure 1). The highest proportion of cancerous mussels was found at site P20 (Croisic Pontoon), and the next highest in the vicinity of this site (P19, P21, and P23; Figure 1).

**Figure 1:**
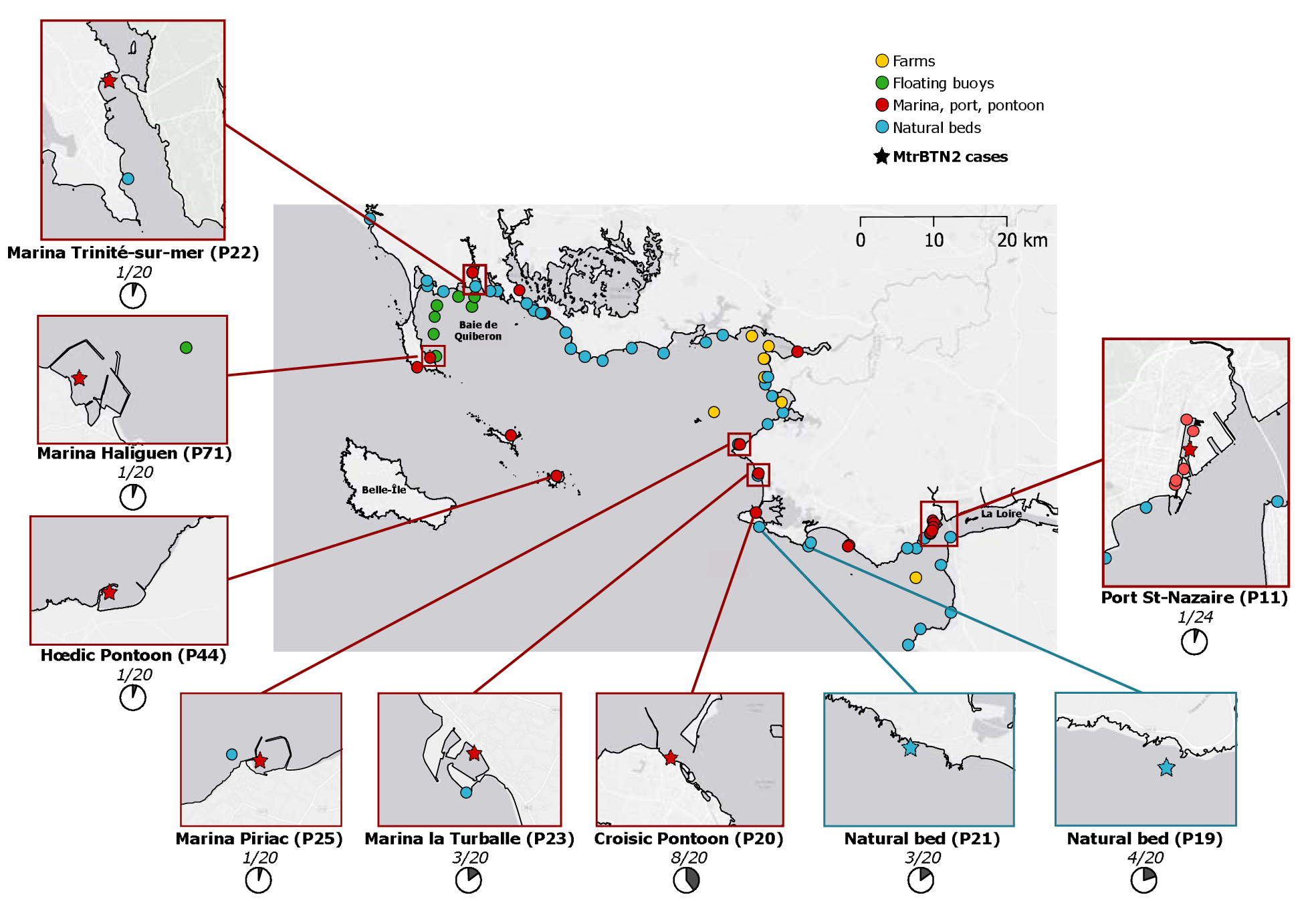
Location of sampling sites. Zoomed-in sites correspond to sites where mussels are affected by MtrBTN2. Pie graphs and numbers in italics correspond to the proportion of cancerous samples in each site. Coloured points represent the type of site, as indicated in the figure’s legend. Map origin: Ersi gray (light).

### M. edulis and M. galloprovincialis are equally affected by MtrBTN2

To determine whether host genetic background or population composition influences cancer prevalence and distribution, mussels were genotyped using 10 SNPs known to be diagnostic between *M. edulis* and *M. galloprovincialis* (the two species coexisting in the studied area).

To reduce the cost of genotyping, we developed a multiplex assay, mixing 10 SNPs that directly give the average ancestry of the sample (G ancestry, Figure S1). This approach was validated by comparing the multiplex fluorescence and average simplex fluorescence of SNPs from a set of samples (Figure S6a). We genotyped each individual from the 9 sites affected by MtrBTN2 (180 mussels) and estimated the average ancestry of each site by genotyping pools of individuals (76 pools). The positive correlation between the average individual fluorescence of a site and the fluorescence of the corresponding pool validated our approach (Figure S6b). Mussels with G ancestry values below 0.3 were assigned to *M. edulis,* those with values above 0.59 to *M. galloprovincialis*, and those in between were assigned as hybrids (Figure 2, FigureS7). Of the mussels affected by MtrBTN2, 15/23 were assigned to *M. edulis*, 7/23 to *M. galloprovincialis* and 1/23 was a hybrid (Table S11). Considering the sites affected by MtrBTN2, MtrBTN2 prevalence did not appear to be significantly different between species with 15/105 (14%) in *M. edulis*, 7/60 (11%) in *M. galloprovincialis* and 1/15 (6%) in hybrids (Fisher’s exact test, p-value = 0.83). Most *M. edulis* mussels affected by MtrBTN2 were found in populations with a majority of *M. edulis* (P44, P23, P20), only two were found in populations with some hybrids and *M. galloprovincialis* mussels (P25, P71) and only one in a population with a majority of *M. galloprovincialis* (P11, Figure 2, Figure S7). Interestingly, all *M. galloprovincialis* mussels affected by MtrBTN2 were found in *M. galloprovincialis* populations (P19, P21, Figure 2, FigureS7), which are close to the site with the highest prevalence (Croisic pontoon, Figure 1). Finally, the hybrid individual affected by MtrBTN2 was found in a population where the two species coexist (P22, Figure 2, FigureS7).

**Figure 2:**
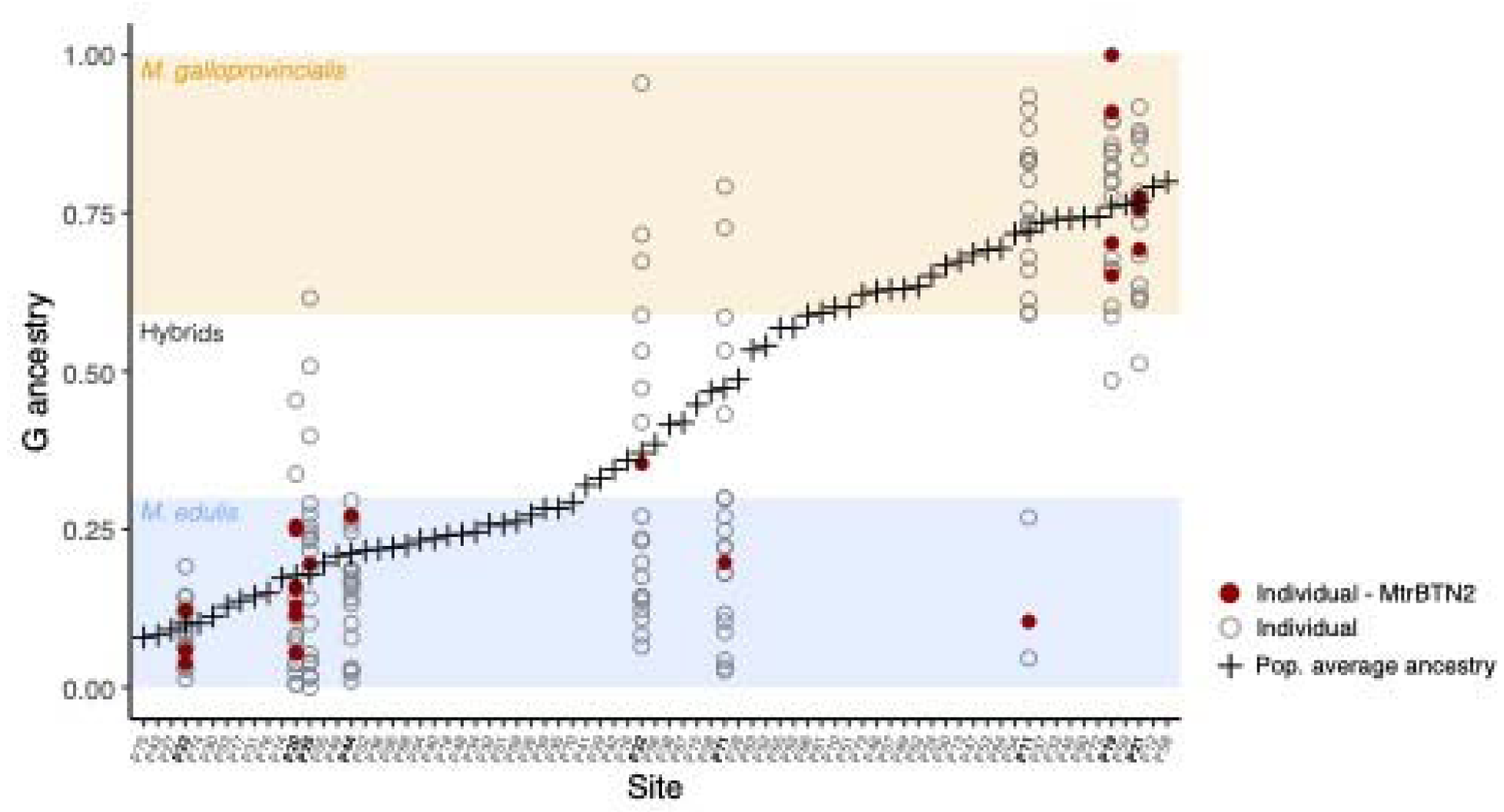
Individual and population ancestry. G ancestry of healthy (empty grey circle) and MtrBTN2 affected (dark red) hosts is shown for the 9 sites with cancer (bold labels). Population average ancestry for all sampled sites is represented by black crosses. Blue, white and orange rectangles correspond to the G ancestry range of *M. edulis*, hybrids and *M. galloprovincialis* respectively.

### 3.1. MtrBTN2 infected mussels are mostly found in ports

After the MtrBTN2 detection step, which revealed 23/1516 individuals infected with this transmissible cancer, we investigated associations with habitat types. Cancerous samples were found in 7 ports (16/23 infected mussels) and 2 natural beds (7/23) near the Croisic pontoon (P20, Figure 1). MtrBTN2 was not detected in the 180 individuals from the 9 mussel farms or in the 140 individuals from the 7 floating buoys. Mussels affected by MtrBTN2 were significantly more frequent in ports than in other habitats (Fisher’s exact test, p-value = 0.0001, Table S12). In addition, the sampled ports are not significantly spatially autocorrelated (Moran I test, p-value = 0.25, Table S13).

### 3.2. Exploring the effect of some environmental variables, none other than the port habitat explains MtrBTN2 prevalence after accounting for spatial autocorrelation

Given that MtrBTN2 was mostly found in ports, we aimed to gain a deeper understanding of whether environmental variables (including population ancestry) could explain this pattern. Univariate correlation between MtrBTN2 prevalence and all environmental variables across habitats are presented in Figure S8. The two best correlations are found with population density and maritime traffic, but we also have strong multicollinearity among variables that needs to be accounted for (see Table S14). The PCA reveals an association between environmental variables and habitat types rather than the presence of cancer itself (Figure 3). Indeed, anthropogenic habitats (ports, floating buoys and farms) with high population density and sheltered from wave action are well separated from natural beds on the second axis of the PCA. Interestingly, maritime traffic (i.e., annual mean of passenger, fishing, and recreational boat density) explains a direction of variation towards human-altered habitats, but also towards sites affected by MtrBTN2. MtrBTN2 seems to be found in sites with buffered temperature (low annual variance) and higher mean annual salinity, environmental variables that are associated with ports in our study. Water temperature indeed tends to be buffered in ports, being warmer in winter and cooler in summer (Figure S4). However, these variables are difficult to interpret as they also explain well the inertia of axis 1 of the PCA which differentiates the sites located near estuaries (La Loire or La Vilaine) from the others. To investigate more rigorously whether some environmental variables could explain prevalence of MtrBTN2, we used a spatial logistic regression model.

**Figure 3:**
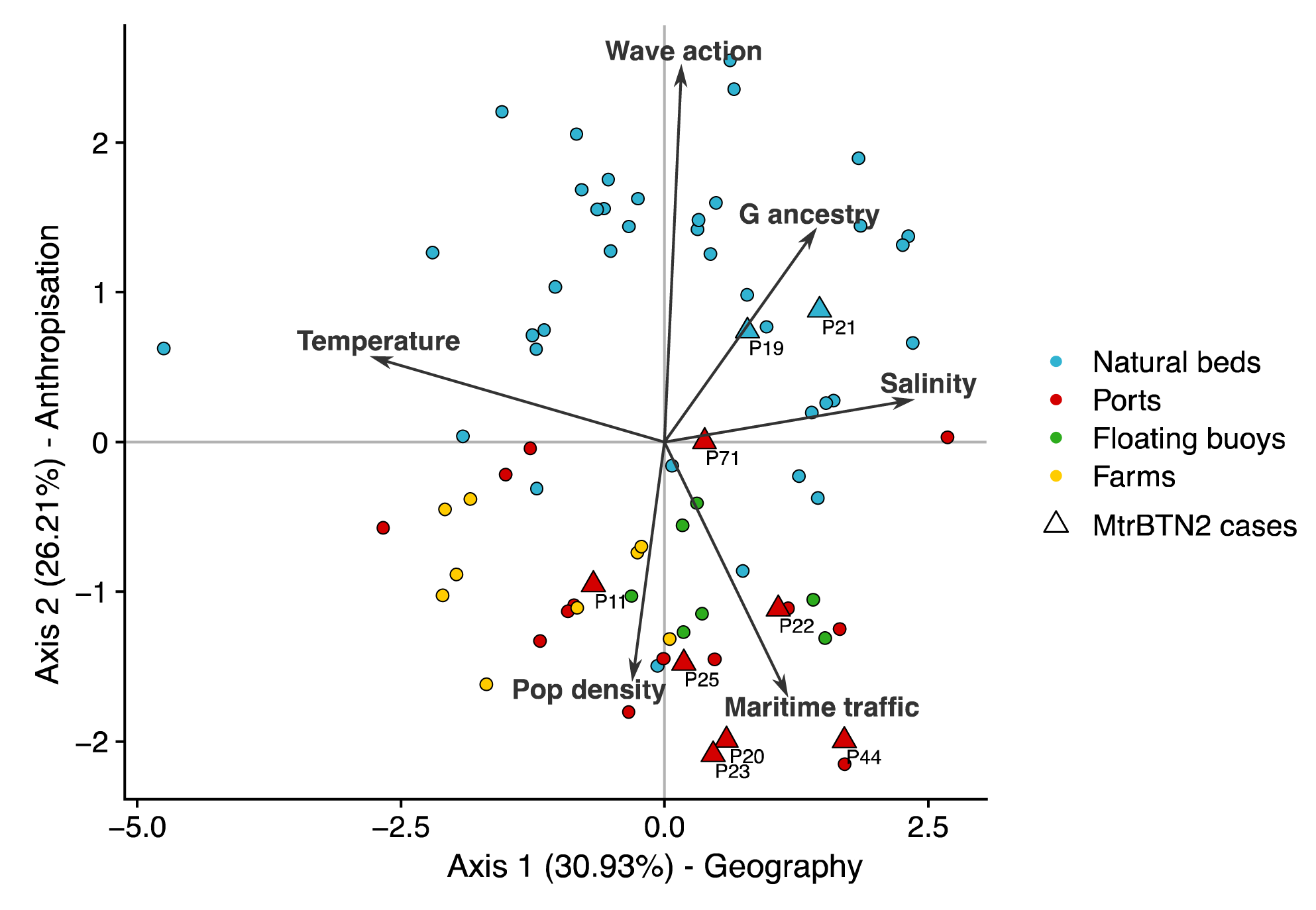
Principal Component Analysis on environmental variables. Axis 1 mainly captures the variance associated with annual temperature variance and annual salinity mean and Axis 2 mainly captures the variance associated with wave action, population density and maritime traffic.

As MtrBTN2 was mostly found in ports (see section 3.4), we included an environmental variable “port” (1 for port habitat, 0 for others) in the regression model analysis. To test for spatial structure, we conducted Moran’s I tests on all environmental variables. The results indicate significant spatial autocorrelation in most environmental variables (Table S13), which was expected as we selected this area due to its mosaic of contrasted environments. However, “population density” and “port” variables were found randomly distributed in space (Table S13). To overcome spatial autocorrelation bias we performed a logistic regression model that corrects for spatial effect (*spaMM* package, [43]) and assessed environmental variables significance via bootstrapped Likelihood Ratio (LR) tests. Bootstrapped LRTs are expected to handle the problem of variables multicollinearity. MtrBTN2 prevalence is significantly explained by the port habitat variable (LR test: χ2L= 7.21, dfL=L1, p-valueL=L0.0072; Table S15), and none of the other environmental variables significantly explain MtrBTN2 prevalence after accounting for spatial effects (Table S15). Despite the residuals following the distributional assumptions of the model (Figure S9) and the significance of the ‘Port’ variable (also supported by the Fisher’s exact test, see 3.4), we acknowledge that quantifying the magnitude of this trend remains challenging and requires further investigations.

## 4. Discussion

Our results show that MtrBTN2 is present at low prevalence along the French Atlantic coast (23/1516 = 1.5%). Strikingly, it is mostly found in ports and is equally distributed in *M. edulis* and *M. galloprovincialis*. Our analysis shows that ports are likely epidemiological hubs for MtrBTN2 propagation. However, the precise environmental factors influencing MtrBTN2 epidemiology have yet to be fully characterised. In this context, we discuss various environmental variables and assess their relevance to the port environment and marine parasite epidemiology.

Most MtrBTN2-infected mussels were found in ports and a prevalence hotspot was found around the Croisic pontoon. Given the low prevalence of MtrBTN2 previously described [16, 32] and confirmed in this study, we acknowledge that our sample sizes per site may be too small (n = 20) to assess MtrBTN2 prevalence with confidence at a site level. However, this was not the scale of resolution we were interested in. Instead, our overall sampling design, capturing replications of habitat features, was chosen to evaluate the effect of habitats and genetic composition of host populations. This allowed us to reveal similar MtrBTN2 prevalence in both species of the mosaic hybrid zone, and a higher prevalence in ports.

The highest prevalence of MtrBTN2 in ports could be explained by two non-exclusive hypotheses. Firstly, ports could offer a favourable environment as it is conducive to disease development and propagation. Indeed, ports are often polluted, confined and permanently immersed habitats (without tidal constraints) with high density of mussels. It is important to note here that our objective is not to explain the emergence of a transmissible cancer, i.e. the initial carcinogenesis in the founder host, but its maintenance and spread. The emergence probably took place long before the industrial period and, despite studying a cancer, the mutagenic effect of pollutants does not have to be put forward to explain our results, but rather their effect on host resistance to infection by transmissible cancer cells. Secondly, there is increased connectivity between ports due to maritime traffic and ports are connected to other sites (other ports or wild sites) through biofouling from vessels.

Ports are often confined areas with longer particle residence times which could increase the contact time between parasites and hosts [46, 47], thereby increasing the probability of transmission and favouring the persistence of a passively dispersing parasite. However, the possible hotspot on Croisic Pontoon, which is affected by strong tidal currents, provides evidence against this hypothesis given that it is not a confined area.

Host population density is an important epidemiological parameter, as high density is known to favour parasite persistence in host populations [48, 49]. Our results suggest that ports are prone to harbouring high population density (Figure 3). However, we found no evidence of MtrBTN2 cases in mussel farms or floating buoys despite high mussel population density in these sites as well. Although population density could favour MtrBTN2 persistence in an affected population, this parameter alone cannot explain the higher prevalence of MtrBTN2 in ports.

Port-based shipping activities are major contributors to water pollution, including issues such as oil spills, plastic contamination, wastewater discharge, and the use of antifouling paint [50, 51]. Pollution is a major environmental stressor that could affect host physiology and increase susceptibility to parasites, particularly by altering its immune response (reviewed in [52]). As filter-feeders, species of bivalve molluscs are particularly concerned [53–55]. However, in our study, most of the affected individuals (15/23) were found in three open sea sites known to be healthy and protected ecosystems with abundant resources (Croisic Pontoon, Natural beds 19 and 21 part of Natura 2000 area; Figure 1), suggesting that pollution alone cannot explain the higher prevalence of MtrBTN2 cancers in ports. Conversely, it can be hypothesised that a healthy, less stressful environment may increase the host permissiveness and allow MtrBTN2 (and possibly other BTNs) to persist at a higher detectable prevalence. Host nutrition can affect disease outcome by driving host immunity and parasite resource availability [56–58]. In invertebrates, increased host nutrition can lead to increased parasite load [59] and improved immunity against parasites [57, 58]. This balance could apply to mussel-MtrBTN2 interaction, and a healthy environment with abundant resources could increase host carrying capacity and/or slow down disease progression. A polluted environment could also affect the survival of MtrBTN2 cells in seawater and reduce their chances of infecting a new host. However, the impact of pollution on mussel hosts and cancer cells needs to be further investigated to clearly establish its influence on the transmission and persistence of such transmissible cancers.

Connectivity through maritime traffic appears to be the main common feature between affected ports, although this explanatory variable does not withstand spatial logistic regression analysis any better than others. Vessels could facilitate MtrBTN2 propagation through transport of affected mussels fixed on the hull. Maritime traffic has been identified as a vector of marine mollusk pathogens (for review see [60]), microorganisms [61] and non-indigenous species [62, 63], through ballast water and/or hull biofouling. As such, ports are considered hotspots of marine non-indigenous species introductions [62, 64, 65] and of marine parasites [66]. *Mytilus* mussels are commonly reported as principal taxa composing hull biofouling (fixed adult mussels; [67–69]). If a port is affected by a local outbreak, it could become a source of contamination to other ports through biofouling of vessels, while natural beds will probably remain isolated by density troughs that could halt the propagation (Figure 4). It is also conceivable that prevalence hotspots exist at other unsampled locations along the coast, either due to environmental factors that have yet to be characterised, or simply due to stochastic local outbreaks. These local outbreaks can be the source of contamination to the anthropogenic connectivity network providing the disease reaches a port. While this does not disqualify the factors discussed above – e.g., pollution, population density and confinement- as drivers of contamination within ports, our study suggests that maritime routes between ports could be anthropogenic gateways to MtrBTN2 propagation. Furthermore, despite a high density of maritime traffic around the floating buoys sampled, these sites remained free of MtrBTN2. Long mooring times in ports increase the probability of hull surface colonisation [69–71] as well as the risk of pathogen transmission [60]. This suggests that, in addition to cruising, ships may have to dock for the spread of MtrBTN2 to be effective, either through transmission of cancer cells or detachment of infected mussels (Figure 4). This could provide an explanation as to why maritime traffic is not a sufficient explanatory variable for MtrBTN2 prevalence. Further investigations would be necessary to ascertain the requisite propagule density (comprising both numerical and time-scale components) for effective transmission, as well as to evaluate possible disruption of byssus quality in affected hosts.

**Figure 4:**
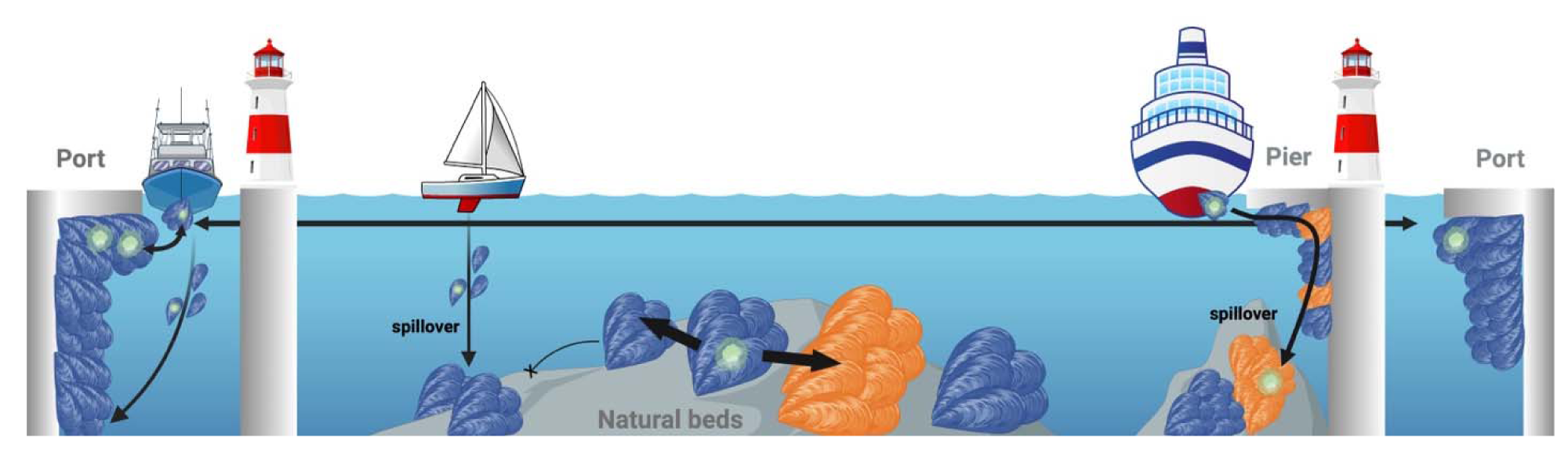
Diagram illustrating the discussion on the propagation of MtrBTN2 across *Mytilus* sp. populations. Arrows’ widths correspond to the probability of MtrBTN2 propagation. Blue and orange mussels correspond to *M. edulis* and *M. galloprovincialis*, respectively. Diagram created with BioRender.

A possible hotspot of MtrBTN2 prevalence was found in the area around the Croisic pontoon. Even though this possible hotspot will need to be confirmed, this site could represent a source for other ports and surrounding natural beds. Indeed, this pontoon is frequently used by maritime shuttles, as well as by recreational and fishing boats, and is connected by maritime traffic to many ports in the studied area. In particular, this pontoon is the departure site of boats travelling to Hoedic (site P44 in Figure 1), an isolated island for which the presence of cancer is most likely explained by anthropogenic transport given the distance from the mainland. MtrBTN2 is also found in foreshores near Croisic. This reveals a possible spillover from ports to adjacent wild populations (Figure 4). Maritime traffic has been identified as a vector for non-indigenous species from commercial ports to natural populations via marinas [72, 73]. In our study, when sites outside standard enclosed ports (e.g., enclosed marinas) were sampled, MtrBTN2 was never detected, suggesting that contagion often remains confined to the port area. Spillover may therefore depend on the degree of confinement of a port.

*Mytilus galloprovincialis* was thought to be more resistant to MtrBTN2 because of lower prevalence in this species [16], and population patches of *M. galloprovincialis* could also have acted as barriers to disease propagation between patches of *M. edulis*. However, our results suggest that *M. galloprovincialis* population patches close to *M. edulis* populations may also show enhanced prevalence. This does not contradict the idea that *M. galloprovincialis* population patches could impose a slightly higher resistance to MtrBTN2 propagation. Under the hypothesis that natural propagation is hindered by density troughs between population patches, or by population patches of a more resistant species, ports and ship traffic would create anthropogenic gateways that favour connections between patches of host populations (Figure 4). This hypothesis could also suggest that BTNs spread naturally and progressively from one host to another by close contact rather than drifting with the currents, making enhanced R0 by anthropogenic dispersal a threat to host populations.

Our results represent a significant advance in our understanding of MtrBTN2 epidemiology, both by proposing ports as epidemiological hubs and by suggesting that the natural propagation of this transmissible cancer could be naturally halted by density troughs in the distribution of host mussel populations. Detection of MtrBTN2 presence on boat biofouling would provide more direct evidence of its role as a vector. It would also be interesting to investigate whether prevalence hotspots are more commonly located in areas with particular environmental conditions, or whether they are stochastic local outbreaks. Sampling vessel biofouling in international ports linking known MtrBTN2 sites (East and North Europe, North West and South Pacific) and nearby marinas, along with genetic comparison of cancer cells, would provide a better understanding of the global and local transmission routes of this transmissible cancer since its emergence. While MtrBTN2 is not associated with extensive mortality events [30], other pathogens could be translocated and affect wild and farmed mussel populations. Various measures are employed to mitigate biofouling, including antifouling painting, dry docking, in-water cleaning with capture, and ballast water filtration (as discussed in [60]). While biofouling management for international vessel exchanges is established [74–76], regulations for biofouling control in local recreational boats remain to be improved [67, 77]. This is a significant concern, as the involvement of recreational boats in the spread of invasive species and possible associated pathogens is increasingly documented [78–80]. Prioritising the survey of vessels that stay in port for long periods [81] or identifying nodes of busy maritime route [72] could contribute considerably to local and international biofouling management.

## 5. Conclusion

Our study reveals that the prevalence of MtrBTN2 along the South Brittany French coast is low, but that the most affected sites are ports (marinas, commercial ports, and docks). This result suggests that ports are epidemiological hubs and that maritime routes are anthropogenic gateways for the propagation of MtrBTN2. Investigations into the presence of other BTNs in boat biofouling would provide more direct evidence of its role as vector, although mussels are the most common species of bivalve mollusc in biofouling. The spread of marine (micro)parasites and pathogens through vessel biofouling is a legitimate concern, and results such as ours -i.e., on a smaller scale than translocations identified in the literature on marine bioinvasion or by [9]- highlight the critical need for politicy regulation to limit the effects of biofouling, both on ship hulls and port docks, in addition to ballast water control.

## Data accessibility

All data used in this study are available in the supplementary material.

## Author’s contribution

Conceptualization, Project administration, Funding acquisition, Writing – review and editing. B., Conceptualization, Funding acquisition, Formal analysis, Investigation, Methodology, Project administration, Supervision, Writing – original draft, Writing – review and editing.

## Acknowledgement

We are grateful to the IAGE company for carrying out the ddPCR assay of this study. We thank the GENSEQ and QPCR HAUT-DEBIT platforms for access to equipments and their expertise. Credit is owed to Olivier Gimenez, François Rousset and an anonymous reviewer for their valuable advice on statistical analysis. Finally, we thank Cécile Perrin for her valuable reading of the manuscript.

---

Supplementary material

Supplementary Tables ( <u>See excel TableS1-15</u>)

**Figure S1:**
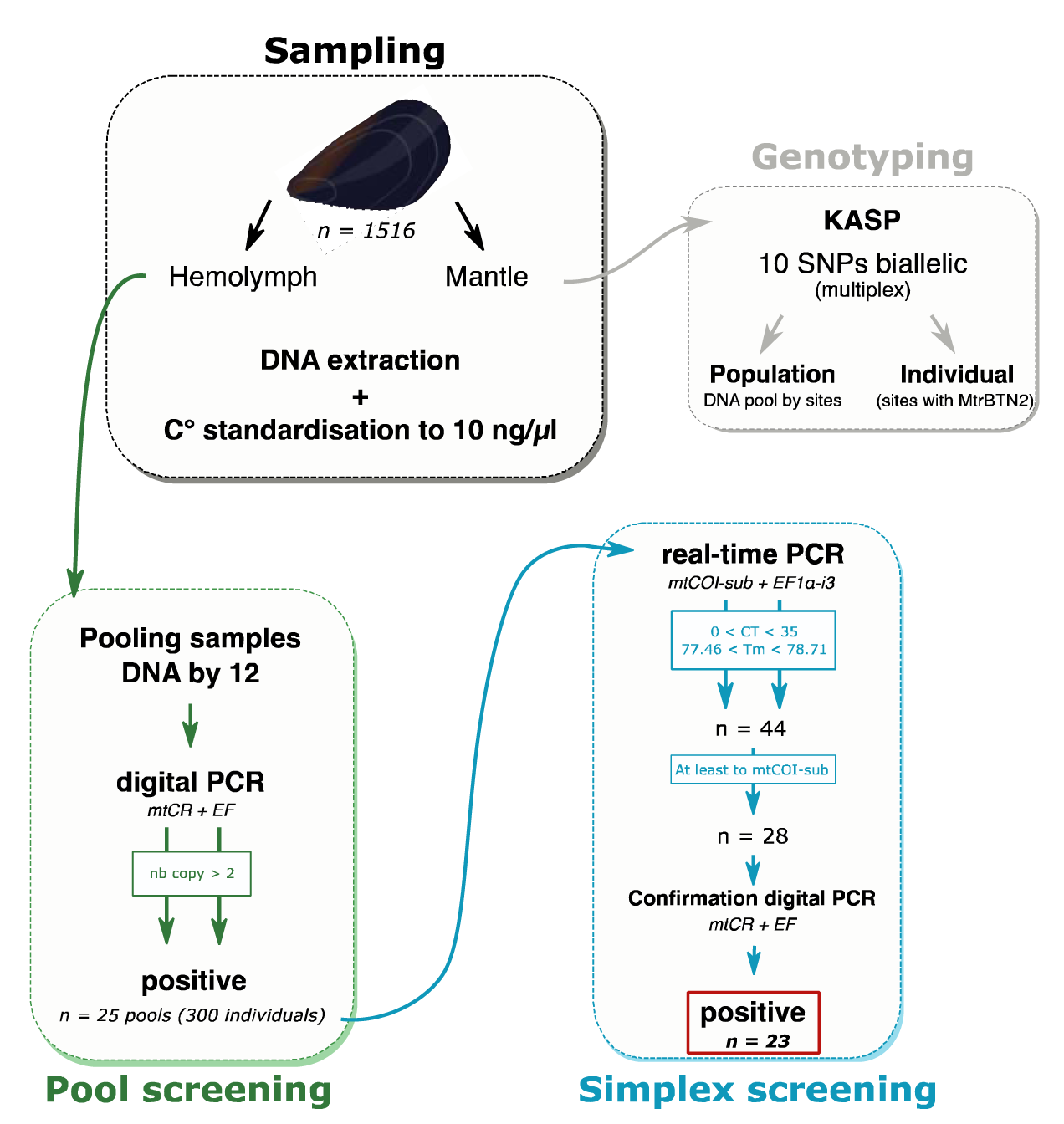
Diagram of screening and genotyping experimental steps.

**Figure S2:**
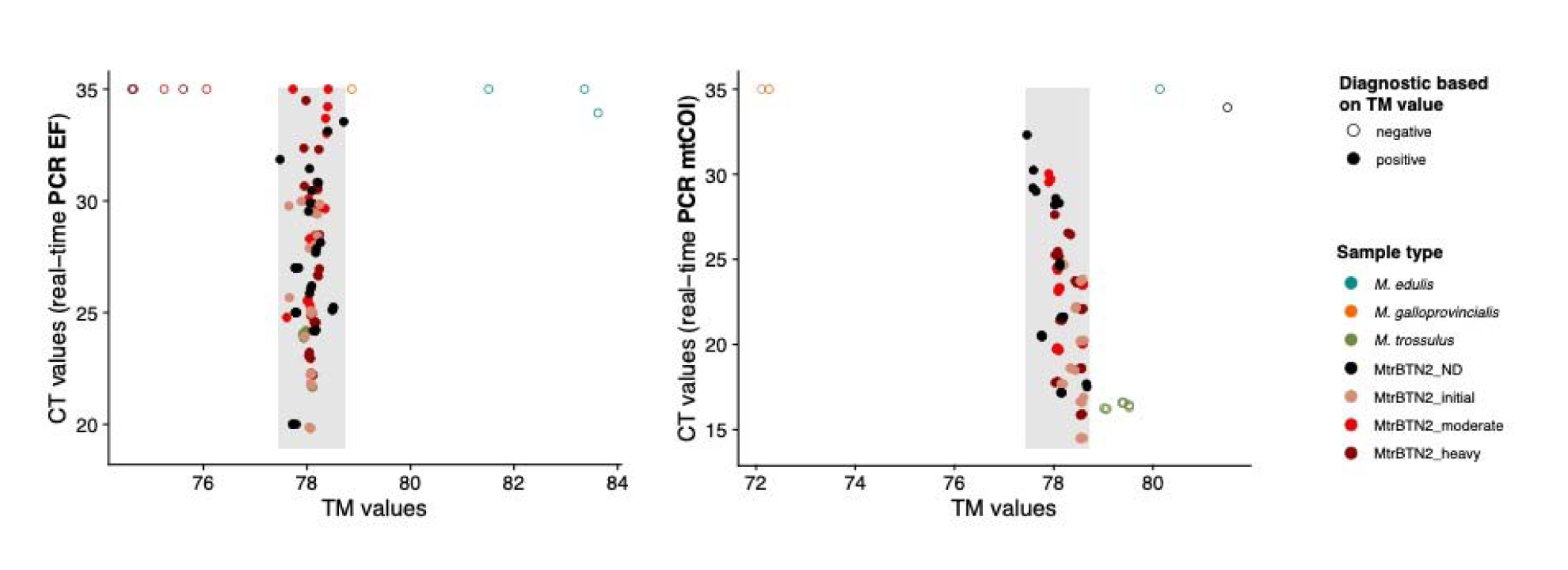
TM threshold determination for EF1α-i3 (left) and mtCOI-sub (right) real-time PCR markers. Grey rectangles correspond to TM threshold based on MtrBTN2 positive control samples (77.46<TM<78.71). Sample type colour and diagnostic based on TM values are indicated in the legend. *M. edulis* and *M. galloprovincialis* correspond to negative controls, *M. trossulus* are *M. trossulus* controls, and MtrBTN2 samples are positive controls (ND: cancer stage not defined, initial: early stage, moderate: moderate stage, heavy: late stage, see TableS3)

**Figure S3:**
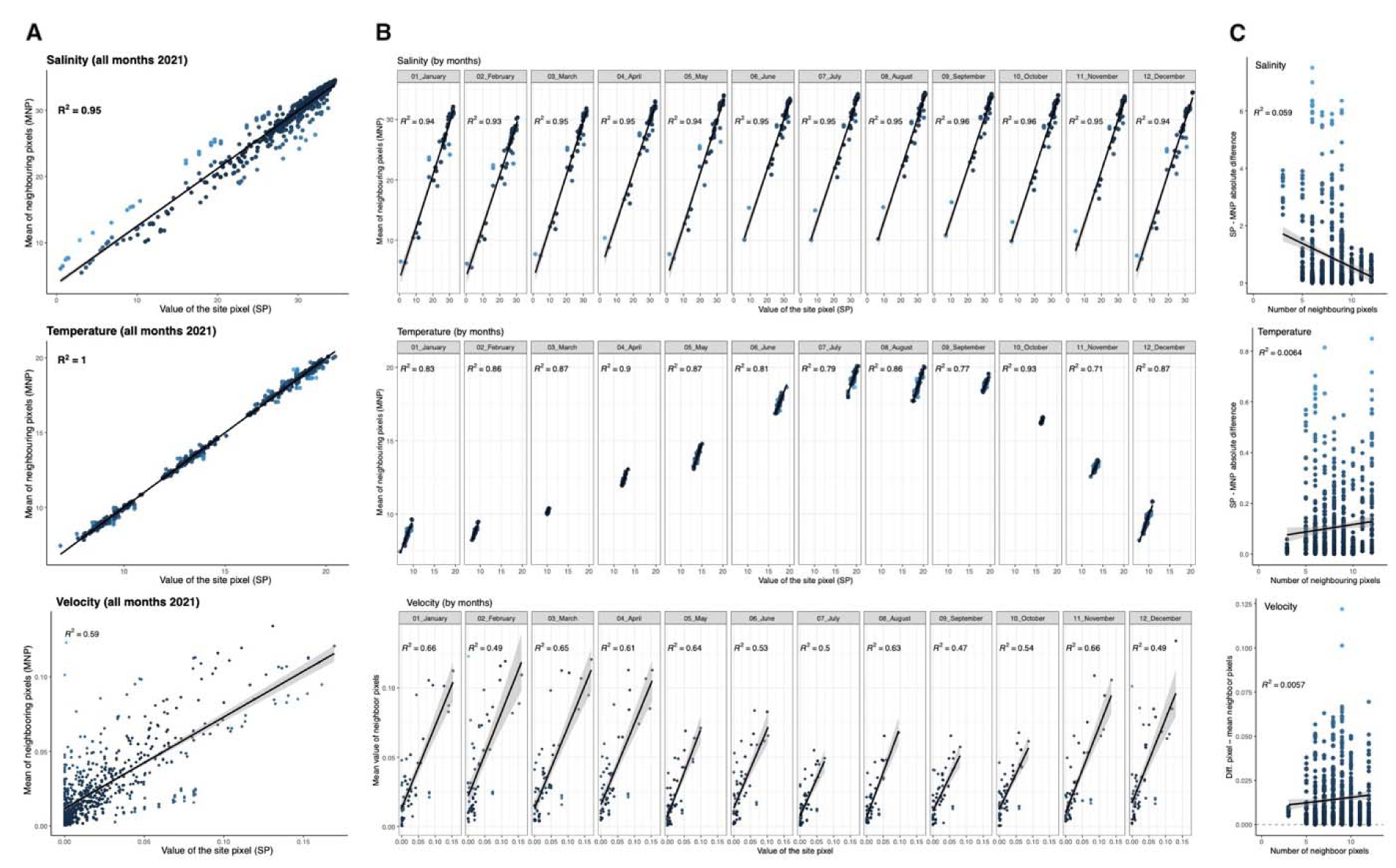
Evaluation of Copernicus model resolution for salinity, temperature and velocity data. Correlation between the value of the site pixel (SP) and the mean value of neighbouring pixels (MNP) for each variable. **(A)** Correlation including all months of year 2021 and **(B)** each month separately. No correlation observed between the absolute difference of SP and MNP values and the number of neighbouring pixels **(C)**.

**Figure S4:**
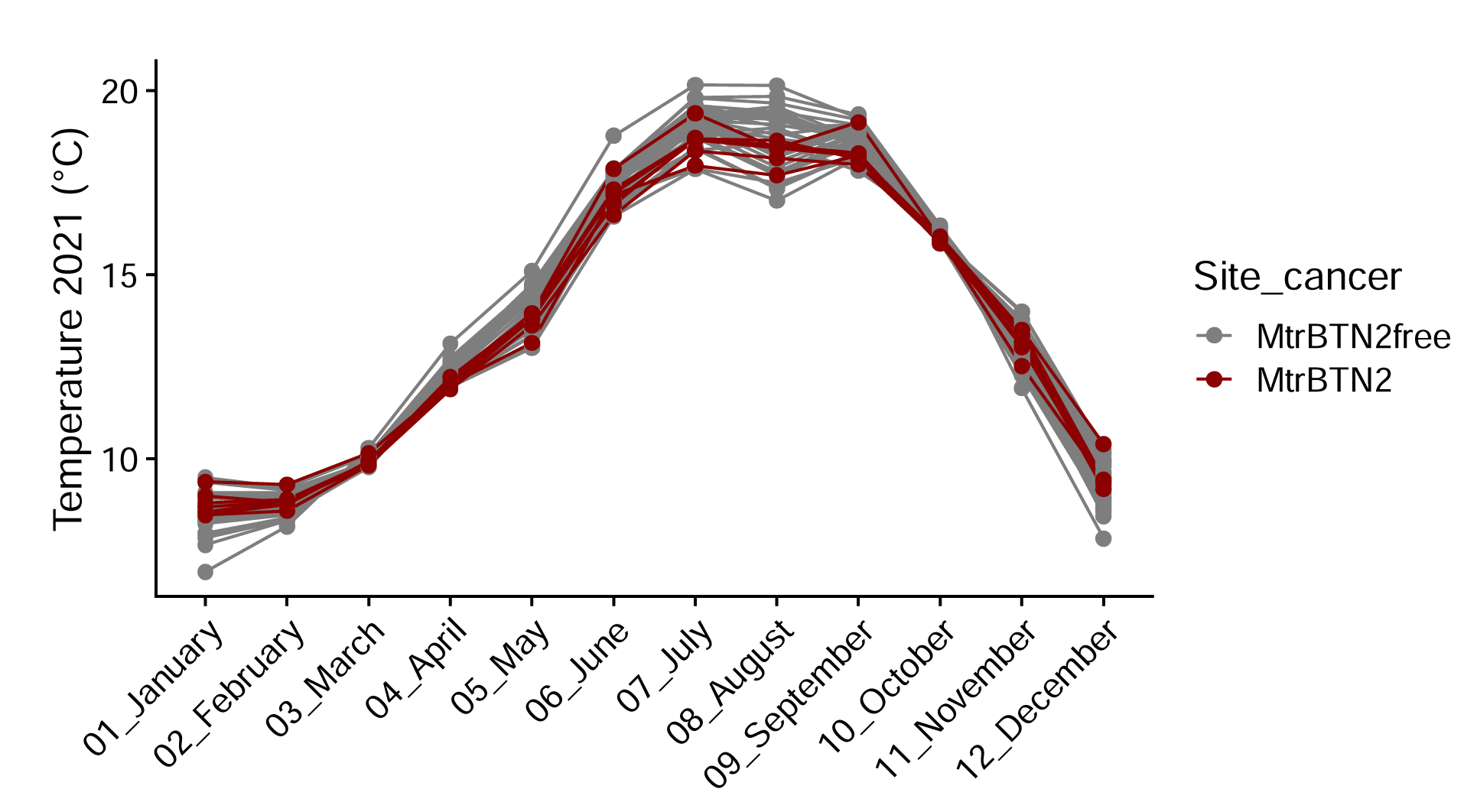
Sampled sites temperature, recorded monthly in 2021. Points correspond to a temperature measurement, solid lines refer to MtrBTN2-free sites (grey) and MtrBTN2 affected sites (darkred).

**Figure S5:**
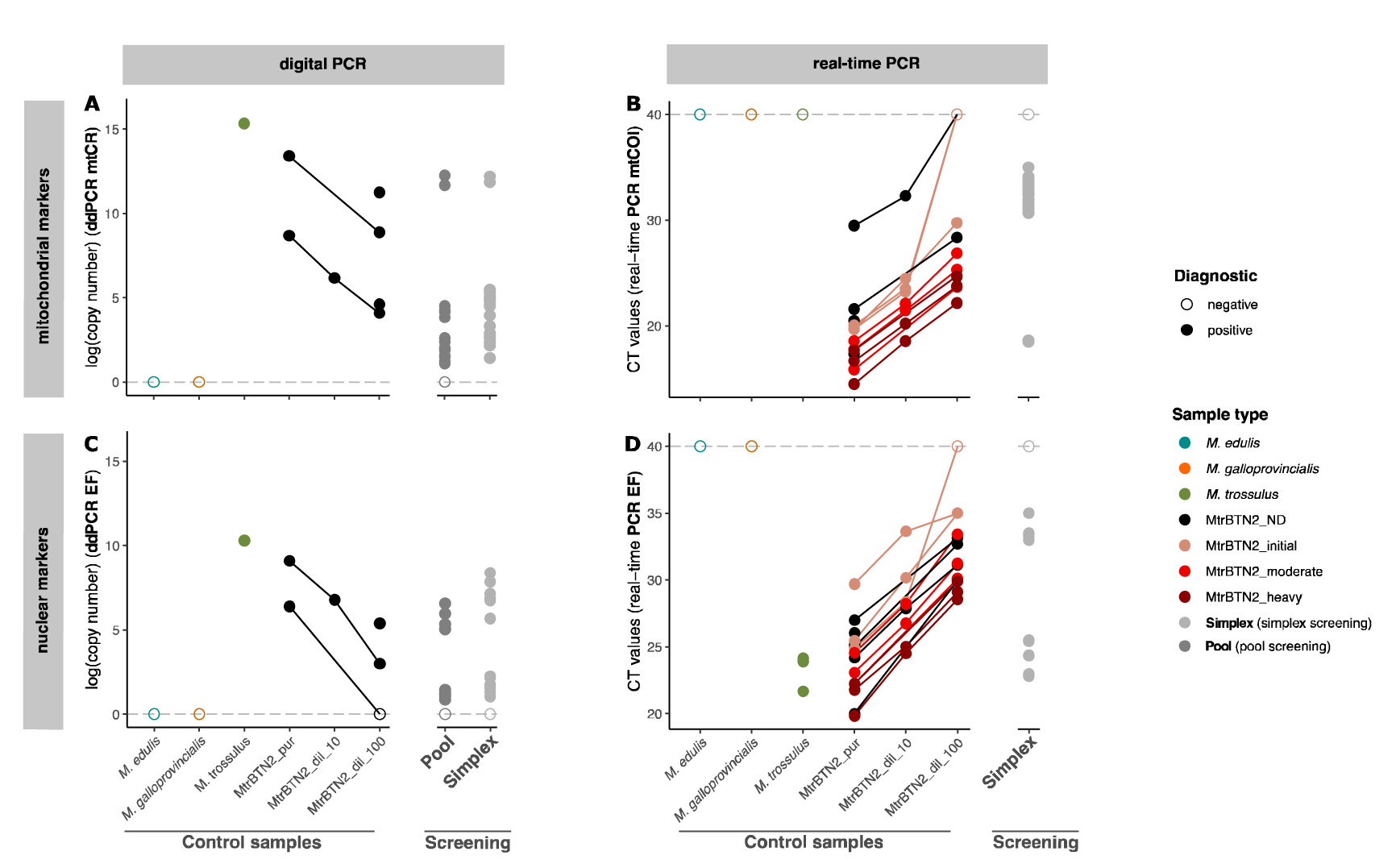
Real-time PCR and digital PCR results. (A) and (C) represent ddPCR results of EF and mtCR, and (B) and (D) represent real-time PCR results of EF1α-i3 and mtCOI-sub, respectively. Dotted lines correspond to the negative thresholds, 0 for no DNA copy detected by ddPCR and 40 for no amplification in real-time PCR. Solid lines in each graph refer to the same sample diluted to different concentrations (pure, x10, x100). *M. edulis* and *M. galloprovincialis* correspond to negative controls, *M. trossulus* are *M. trossulus* controls, and MtrBTN2 samples are positive controls (ND: cancer stage not defined, Initial: early stage, moderate: moderate stage, heavy: late stage, see TableS2). Screening results using both method and markers are shown for Pool and Simplex. Sample type colours and diagnostic status are indicated in the figure legend.

**Figure S6:**
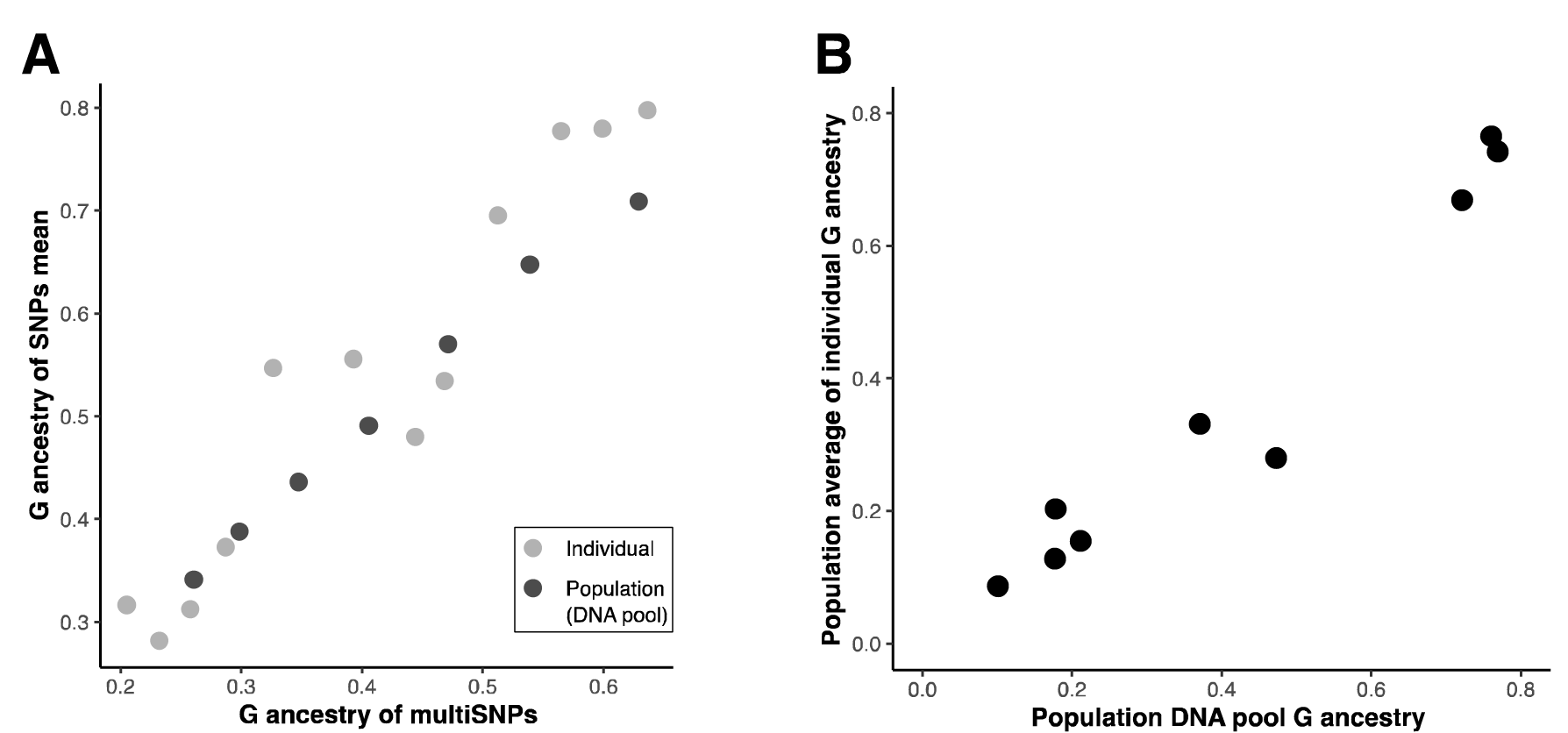
Genotyping method validation. (A) Positive correlation between multiSNPs G ancestry and the mean of the 10 single SNPs from a subset of Individual and Pool samples. **(B)** Positive correlation between population DNA pool G ancestry and the mean individual G ancestry of individuals from the same population.

**Figure S7:**
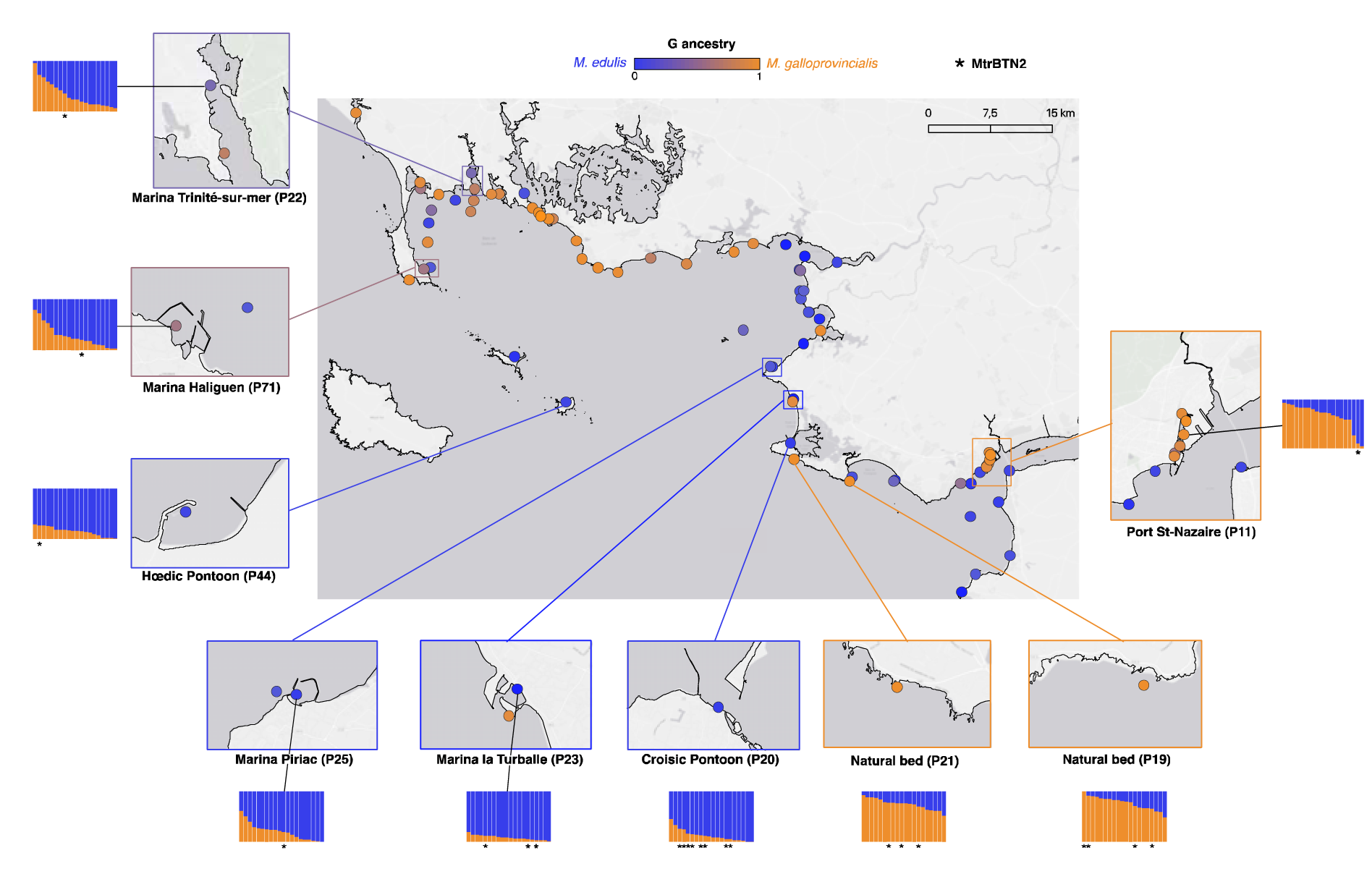
Population and individual ancestry composition across the sampled area. Zoomed-in sites correspond to sites with mussels affected by MtrBTN2. Barplots represent the estimated ancestry of individuals based on the 10SNPs multiplex tool. MtrBTN2 individuals are indicated by stars under each barplot. Coloured points represent the population average ancestry, as indicated in the figure’s legend. Map origin: Ersi gray (light).

**Figure S8:**
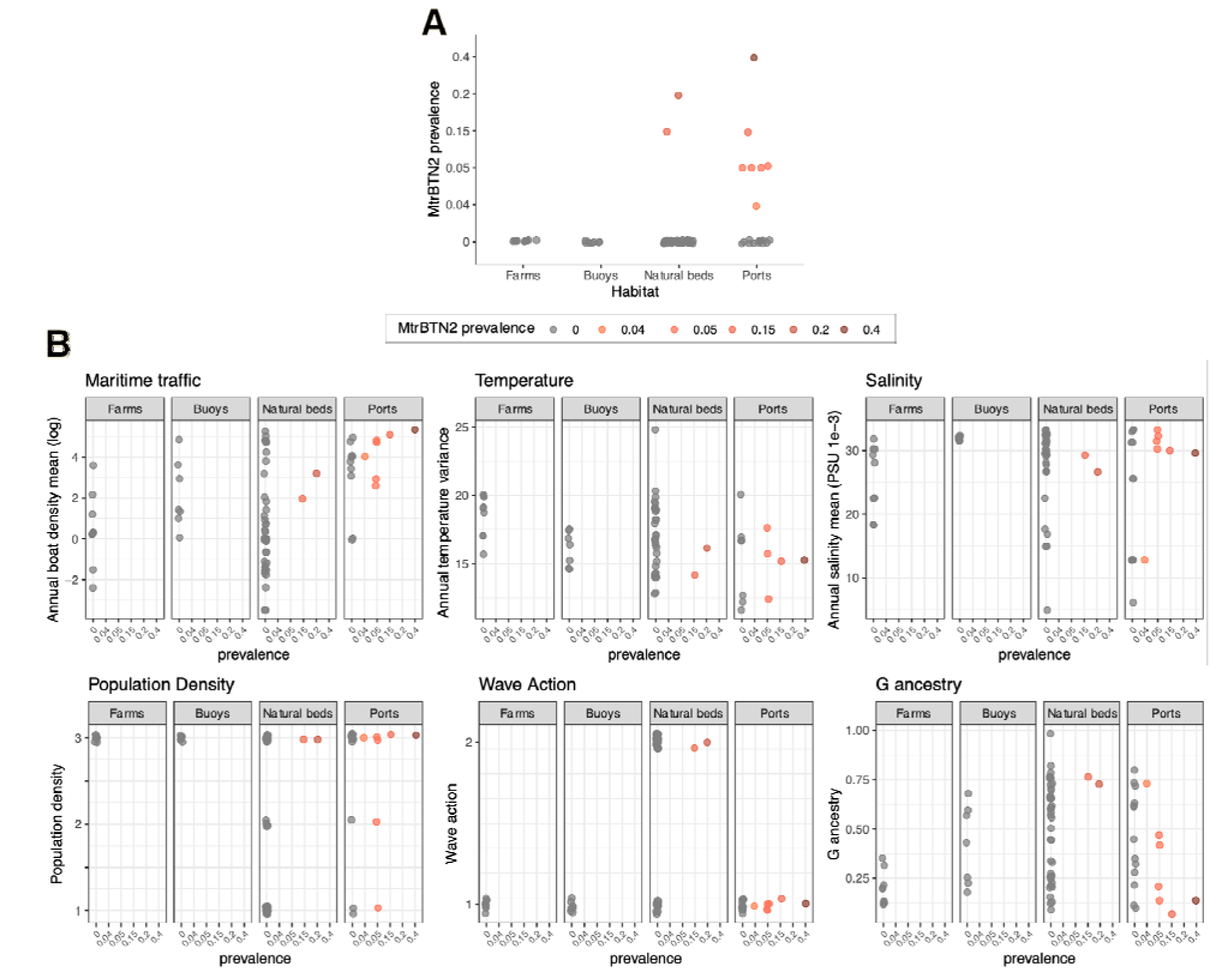
Univariate correlations between the prevalence of MtrBTN2 and environmental variables. (A) MtrBTN2 prevalence among habitats. (B) Correlations for the six environmental variables studied. Coloured points correspond to the prevalence of MtrBTN2 as indicated in the legend.

**Figure S9:**
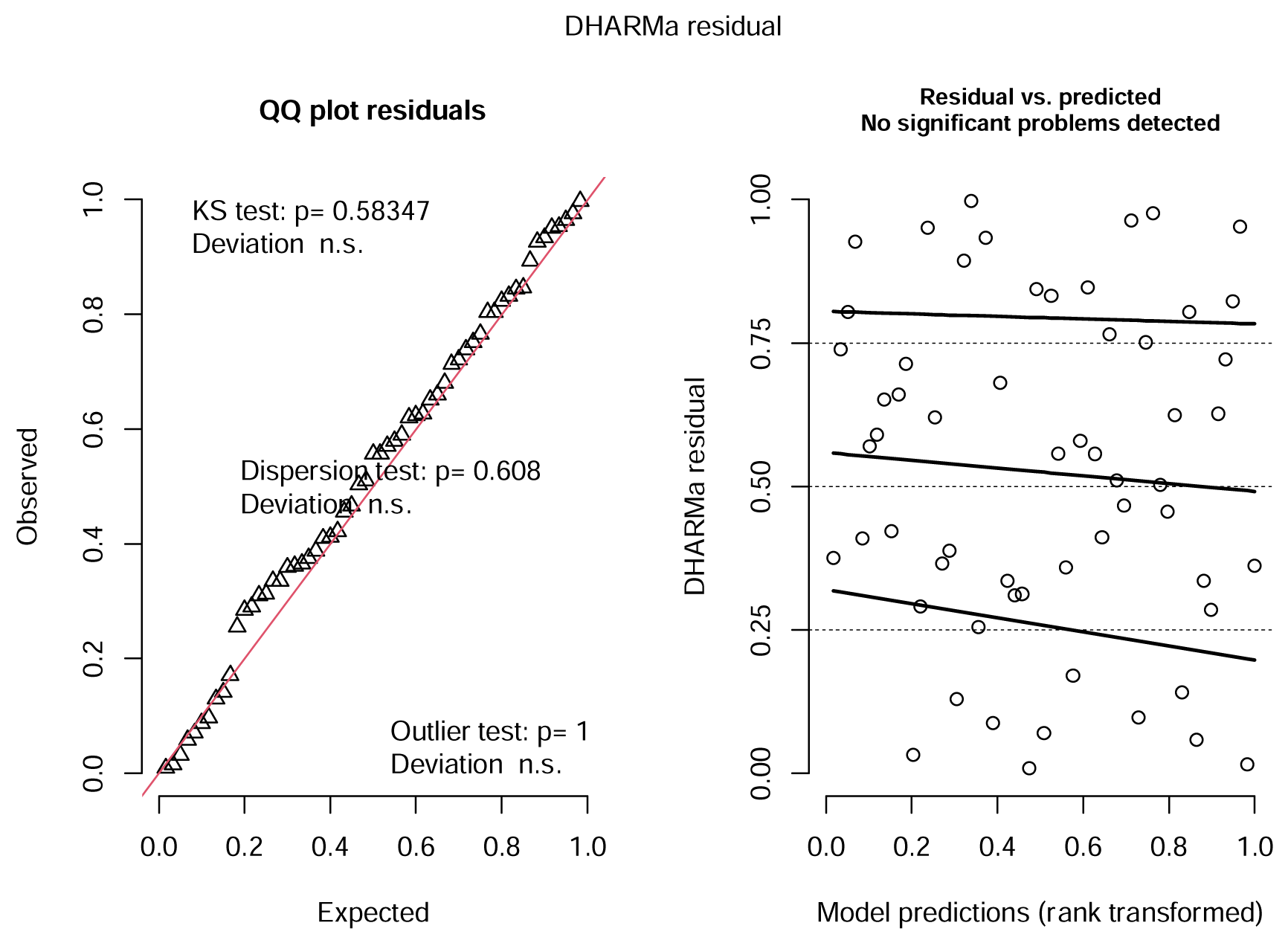
Residual diagnostics of the logistic regression model accounting for spatial autocorrelation. Model results are presented in Table S15. Plot generated using *simulateResiduals()* function of DHARMa package in R.

**TableS1 : Description of sampled sites.** Sources of environmental variables are provided in TableS5. Site: site name; N: number of sampled mussels; MtrBTN2 prevalence: number of MtrBTN2 affected mussels; Temperature: annual variance; Salinity: annual mean; Maritime traffic: annual mean in log; temp: temperature; sal: salinity; vel_E and vel_N: Eastward and Northward current velocity, respectively; [fish-pass-other][19–21]_[1–4] fish: fishing boat density, pass: passenger boat density, other: recreational boat density, 19 to 21 correspond to the year of record, and 1 to 4 correspond to spring, summer, autumn, winter, respectively. Temperature and Salinity missing data (blank) correspond to sites outside of Copernicus model pixels (n = 13).

**Table S2: Primer and probe sequences used in real-time PCR and digital PCR**.

**Table S3: Control samples used to validate real-time PCR and ddPCR screening tools.** All samples are from our laboratory sample collection. *M. edulis* and *M. galloprovincialis* correspond to negative controls, *M. trossulus* are *M. trossulus* controls, and MtrBTN2 samples are positive controls. MtrBTN2 stages were defined by cytology of the hemolymph, according to the following classification: early (<15% cancer cells), moderate (15-75% cancer cells), late (>75% cancer cells), ND when cancer stage was not defined.

**Table S4: Bi-allelic nuclear SNPs used for the multiSNPs genotyping**.

**TableS5: Source and description of environmental variables collected for each habitat (ports, natural beds, farms, buoys)**.

**TableS6: Data for the evaluation of Copernicus model resolution.** Sources of environmental variables are provided in TableS5. Site: site name; N_pixels: number of neighbouring pixels within a buffer zone of 0.05°; SP: value of the site pixel; MNP: mean value of the neighbouring pixels; SP_MNP: absolute difference between SP and MNP values; .s, .t and .v refer to salinity, temperature and velocity values, respectively. Velocity correspond to the combination of squared Northward and Eastward velocity values. Missing data (blank) correspond to sites outside of Copernicus model pixels (n = 13).

**Table S7: Control sample results validate real-time PCR and ddPCR screening tools.** EF_CopyNb: copy number for EF ddPCR marker; mtCR_CopyNb: copy number for mtCR ddPCR marker; CT: threshold cycle, TM: melting temperature for EF1a-i3 and mtCOI-sub real-time PCR markers. NP: not performed.

**TableS8: Pooled and simplex screening results.** Pool_CopyNb_EF: copy number for EF ddPCR marker (pooled screening); Pool_CopyNb_mtCR: copy number for mtCR ddPCR marker (pooled screening); CT: threshold cycle, TM: melting temperature for EF1a-i3 and mtCOI-sub real-time PCR markers; Simplex_CopyNb_EF: copy number for EF ddPCR marker (simplex screening); Simplex_CopyNb_mtCR: copy number for mtCR ddPCR marker (simplex screening). NP: not performed.

**TableS9: Simplex screening results for positive samples.** CT: threshold cycle, TM: melting temperature for EF1a-i3 and mtCOI-sub real-time PCR markers; PCR_result: marker name positive in real-time PCR; EF_CopyNb: copy number for EF ddPCR marker; mtCR_CopyNb: copy number for mtCR ddPCR marker; ddPCR_result: marker name positive in ddPCR; MtrBTN2: final diagnostic status. NP: not performed.

**Table S10: Correspondence between pool and simplex screening.** Pool_CopyNb_EF : copy number for EF ddPCR marker (pooled screening); Pool_CopyNb_mtCR : copy number for mtCR ddPCR marker (pooled screening); Pool_ddPCR_result: marker name positive in ddPCR (pool); Nb_ind_positive: number of individuals in the pool found positive for one or two marker in simplex screening; Simplex_PCR_result: marker name of the positive individual in real-time PCR; Simplex_ddPCR_result: marker name of the positive individual in ddPCR, negative if the sample did not amplify. NP: not performed.

**Table S11: MultiSNPs genotyping results.** G ancestry: fluorescence value; Ancestry: ancestry defined based on G ancestry values.

**Table S12: Contingency table used for the Fisher exact test**.

**Table S13: Moran I test on environmental variables.** Results were obtained using moran.test() function in R.

**Table S14: Correlation coefficient and Spearman correlation test results for each environmental data pairs**.

**Table S15: Spatial logistic regression model testing the effect of environmental variables on MtrBTN2 prevalence.** Model results were obtained using fitme() from spaMM package in R. Matern covariance function was included to account for spatial autocorrelation (i.e. Matern(1 | Longitude + Latitude). Effects of environmental variables were tested using Likelihood ratio (LR) tests using fixedLRT() function from the same package. Temperature: annual variance; Salinity: annual mean; Maritime traffic: annual mean.

